# DNA Combing *versus* DNA Spreading and the Separation of Sister Chromatids

**DOI:** 10.1101/2023.05.02.539129

**Authors:** Alice Meroni, Sophie E. Wells, Carmen Fonseca, Arnab Ray Chaudhuri, Keith W. Caldecott, Alessandro Vindigni

## Abstract

DNA combing and DNA spreading are two central approaches for studying DNA replication fork dynamics genome-wide at single-molecule resolution by distributing labeled genomic DNA on coverslips or slides for immunodetection. Perturbations in DNA replication fork dynamics can differentially affect either leading or lagging strand synthesis, for example in instances where replication is blocked by a lesion or obstacle on only one of the two strands. Thus, we sought to investigate whether the DNA combing and/or spreading approaches are suitable for resolving adjacent sister chromatids during DNA replication, thereby enabling the detection of DNA replication dynamics within individual nascent strands. To this end, we developed a thymidine labeling scheme that discriminates between these two possibilities. Our data suggests that DNA combing resolves single chromatids, allowing the detection of strand-specific alterations, whereas DNA spreading does not. These findings have important implications when interpreting DNA replication dynamics from data obtained by these two commonly used techniques.

## Introduction

DNA replication is the key process that ensures cell division and the correct propagation of genetic information to daughter cells. This process entails the transient unwinding of the DNA duplex, leading to the formation of a three-way junction structure within which the separated parental DNA strands serve as a template for the synthesis of the newly formed leading and lagging strand fragments. This process is mediated by a complex of proteins termed the “replisome” (Yao and O’Donnell 2009).

One of the most powerful and widely used techniques to monitor DNA replication fork dynamics at single-molecule resolution is the DNA fiber spreading assay, which relies on the ability of replicating cells to incorporate thymidine analogs in the newly synthesized (nascent) strands. Briefly, nascent strands are sequentially pulse-labelled with two labelled thymidine analogues, typically chosen from 5-iodo-2-deoxyuridine (IdU), 5-chloro-2′-deoxyuridine (CldU), or 5-bromo-2-deoxyuridine (BrdU). Cells are then lysed to release and deposit their DNA on the positively charged surface of a silanized slide. The slide is then tilted at a 25-60 degrees angle to favor the spreading of the DNA by gravity (Parra and Windle 1993). Alternatively, in a variation of DNA spreading known as DNA combing, the labelled cells are embedded in agarose plugs from which DNA is extracted, usually with the aid of proteinase K treatment, and ‘combed’ in uniform parallel arrays with the aid of a combing machine (Bensimon et al. 1994).

After spreading or combing, the labelled DNA is visualized through immunofluorescence by incubation with antibodies that specifically recognize the halogenated nucleosides incorporated in the DNA. The length of individual DNA fibers can be measured at single molecule level as a direct readout of DNA replication progression (Techer et al. 2013). A fiber of approximately 1 μm corresponds to 2.6 kb of DNA with the DNA spreading technique (Daigaku, Davies, and Ulrich 2010; Jackson and Pombo 1998) and approximately 2 kb for DNA combing (Bensimon et al. 1994; Michalet et al. 1997). Alternatively, the incorporation of thymidine analogs can be quantified by mass spectrometry, using the MS-BAND approach (Ashour et al. 2023). In addition to DNA replication fork progression, other replication parameters can be evaluated using the DNA fiber approach including fork symmetry, origin firing, and nascent DNA degradation (reviewed in (Quinet et al. 2017)).

DNA spreading is less time-consuming than DNA combing and is generally regarded as a higher-throughput technique. On the other hand, DNA spreading leads to a non-uniform distribution of the DNA on the slides, whereas DNA combing leads to a parallel and uniform distribution of DNA molecules. This uniform distribution enables more robust and accurate measurements of the length of the DNA fibers, which is important when measuring parameters such as the DNA replication velocity or the distance between two replication origins within the same filament (Techer et al. 2013).

An important consideration is that the thymidine analogs are simultaneously incorporated on both the leading and lagging strands. However, the genome is constantly under the attack of endogenous and exogenous agents that can lead to DNA damage and perturb DNA replication dynamics (Berti, Cortez, and Lopes 2020; Cybulla and Vindigni 2023).

These lesions or replication obstacles can be located in one or both template DNA strands and thereby differentially affect either the leading or the lagging strand synthesis. Here, we combined the efforts of three independent laboratories to evaluate whether DNA spreading and DNA combing assays can resolve distinct DNA sister chromatids. This is important because DNA replication defects in only one of the two sister chromatids, such as a broken sister chromatid arising from DNA replication fork collapse at a single-strand break in one of the two DNA template strands, can only be detected if the two adjacent sister chromatids are separated during the DNA fiber experiment (Fig.1A). We predicted that whilst DNA combing likely results in sister chromatid separation, because of the more ‘aggressive’ lysis conditions and inclusion of proteinase K, DNA spreading might not do so.

**Fig. 1.**
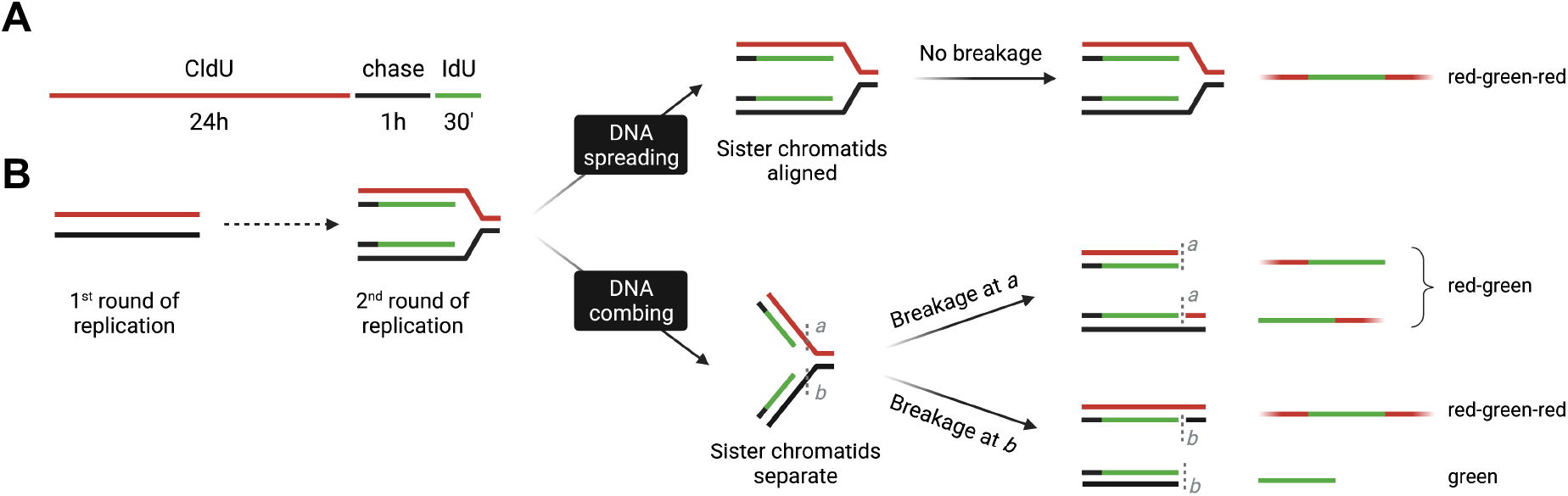
Schematic and model for defining the outcome of DNA spreading and DNA combing on sister chromatid separation. **(A)** Use of halogenated thymidine analogs to distinguish between sister chromatids during S phase. RPE-1 cells are incubated with 1 μM CldU for 24 hours (first round of replication), chased with fresh media for 1 hour, and incubated with 200 μM IdU for 30 minutes (second round of replication). **B)** Predicted outcomes for sister chromatid separation during DNA spreading (top) and DNA combing (bottom). During DNA spreading, we propose that sister chromatids remain aligned/adjacent to each other, resulting in the fiber classifications shown on the right. We propose that because sister chromatids likely remain aligned and/or adjacent to each other, there is little or no mechanical breakage of the replication fork during DNA spreading. In contrast, during DNA combing, we propose that sister chromatids become separated as a result of the more extensive cell lysis conditions and inclusion of proteinase K (which will degrade cohesin). We suggest that during combing, at least one of the ‘splayed’ arms at such DNA replication forks will break within the single-stranded region at either position *a* or *b*, leading to linear fibers with the classifications shown on the right.

## Results

To determine the ability of the spreading and combing assays to separate the two-sister chromatids, we designed a labeling scheme in which we first incubated human retinal pigment epithelial cells (RPE-1) cells with CldU (red label) for 24 h to allow a single round of DNA replication and thus label one DNA strand of the entire genome of all cells with CldU. Next, we incubated the cells for 30 min with IdU (green label) to pulse label active DNA replication forks during the second round of DNA replication. Importantly, we chased cells with media lacking labelled nucleoside for 1 hour between the two analogs to separate CldU from IdU incorporation during the second round of replication (Fig. 1A). We then collected the cells and processed them in parallel with the DNA spreading and combing approaches. We reasoned that if adjacent sister chromatids remain together, we should detect primarily replication tracts with adjacent red and green signals (Fig 1B). In contrast, if sister chromatids are separated, we should also detect green-only tracts. This is because if sister chromatids are separated, the resulting ‘splayed’ forks likely break mechanically during the combing process, resulting in linear fibers comprised of individual sister chromatids. Consequently, we should now be able to detect those nascent DNA replication tracts present on the previously unlabelled template DNA strand (green-only tracts in Fig.1B).

Consistent with our hypothesis, following DNA combing, ∼40% of labelled DNA replication tracts were green-only, with ∼60% being red-green or red-green-red (Fig.2). These data suggest that adjacent sister chromatids are indeed separated during DNA combing, most likely as a result of the more extensive lysis conditions and/or use of proteinase K in this approach, leading to subsequent breakage of the splayed arms of the DNA replication fork during the DNA combing process.

**Fig. 2.**
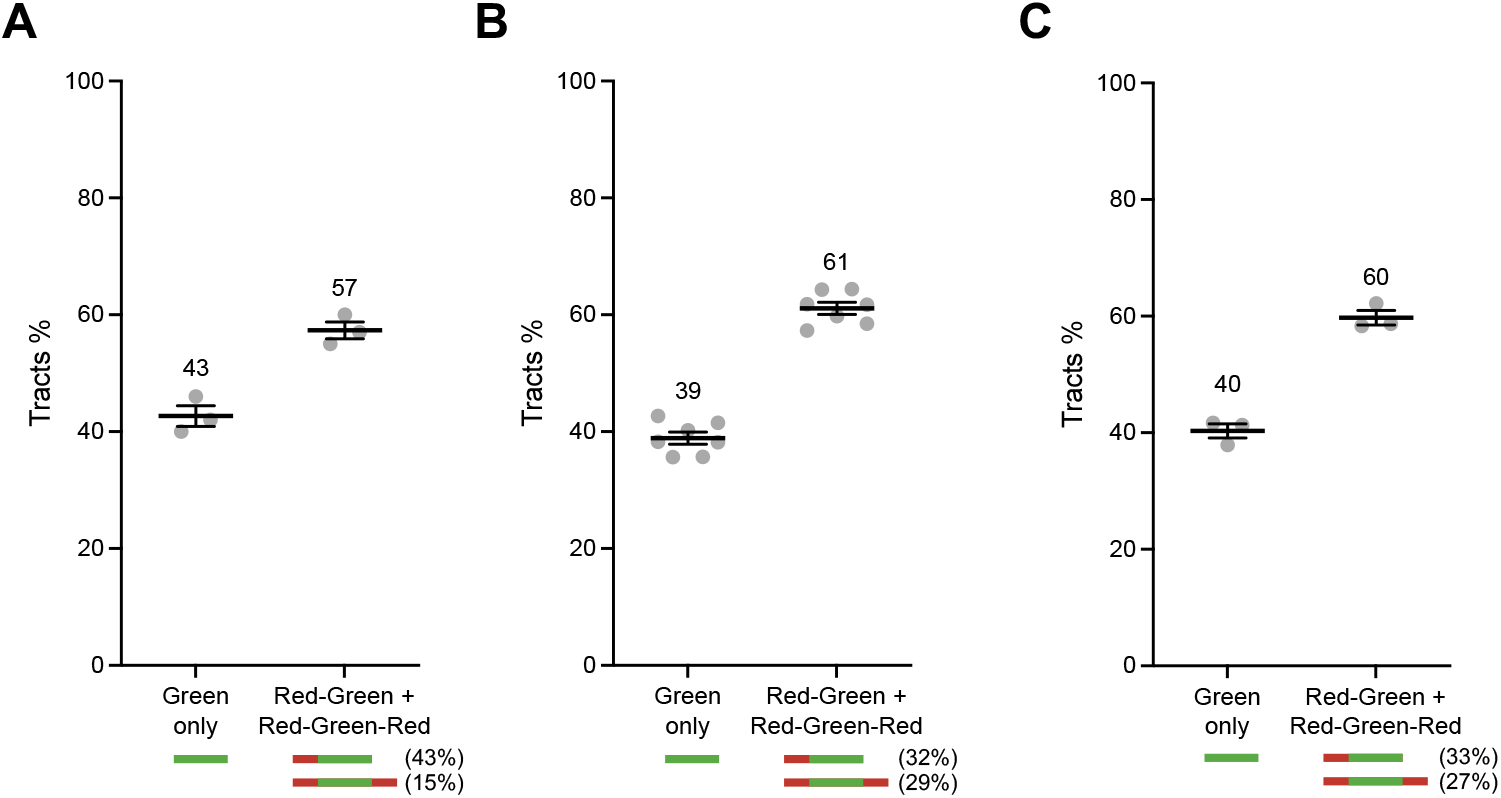
Sister chromatids are separated during DNA combing. **(A-C)** DNA combing assay performed as depicted in Fig. 1A. Green-only *versus* Red-Green or Red-Green-Red tracts are scored and represented as percentage of total pulse-labeled tracts. Numbers indicate the mean tracts %. **A-C** indicate independent data sets from the Vindigni, Caldecott, and Chaudhuri laboratories, respectively (*N*=3 in **A**, *N*=6 in **B, and** *N*=3 in **C**).

To confirm the high level of mechanical fork breakage during DNA combing, we labelled nascent DNA for 30 minutes with IdU followed by staining with anti-IdU and anti-DNA antibody (anti-ssDNA) to detect the integrity of the DNA flanking the DNA replication tract, directly (Fig 3A). We then quantified those pulse-labeled tracts that had DNA on one side only or on both sides and defined these as representing broken sister chromatids and intact sister chromatids, respectively (Fig 3A). We found that ∼50% of DNA replication tracts lacked DNA continuity on one side of the tract, and thus were a result of fork breakage. Since each broken fork results in one broken and one intact (but nicked) sister chromatid, we conclude that most, if not all, DNA replication forks break at a single-stranded junction during DNA combing (Fig 3B).

**Fig. 3.**
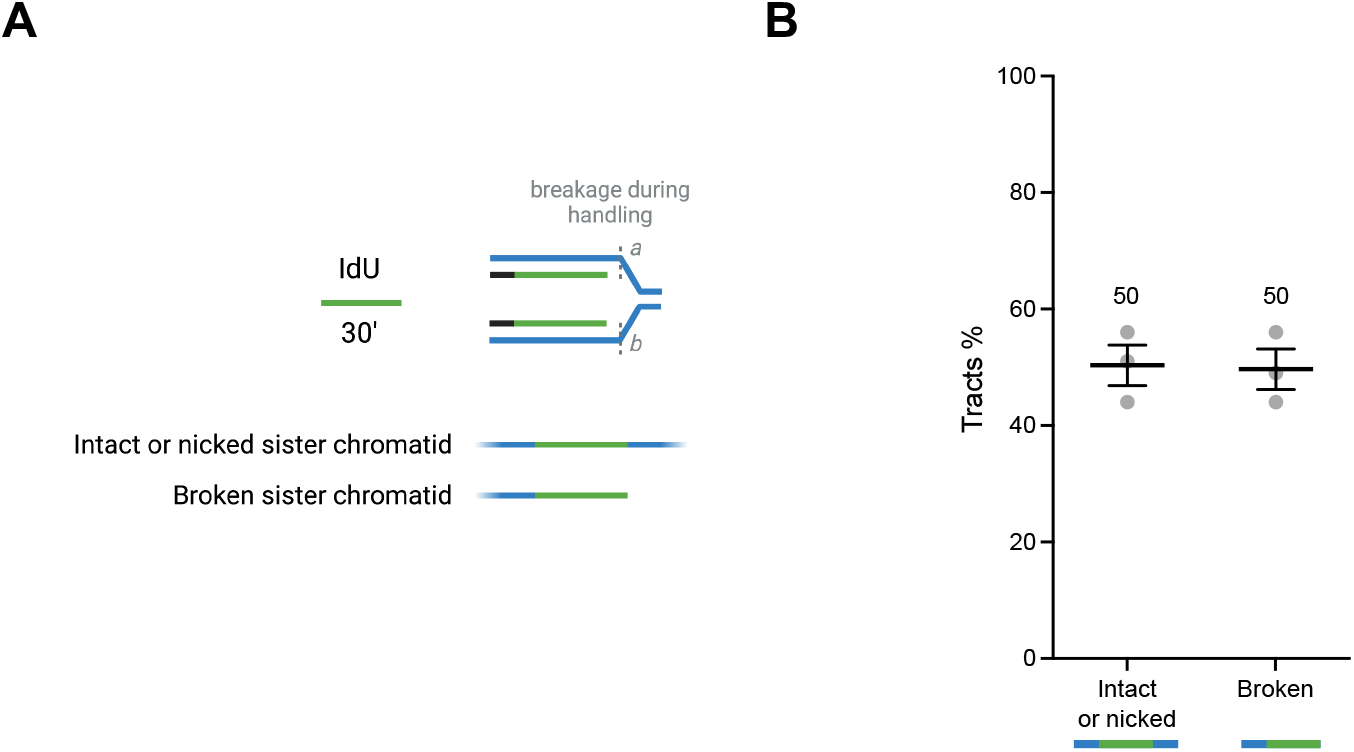
Fork breakage occurs during DNA combing. **(A)**, Schematic describing the use of pulse-labeling to measure fork breakage. RPE-1 cells were incubated with 100 μM IdU for 30 minutes. IdU was detected as above and total DNA was detected using anti-ssDNA antibody. Broken sister chromatids contain DNA on only one side of the IdU pulse label, whereas intact or nicked sister chromatids contain IdU pulse label that is flanked on both sides by DNA (blue). Note that fork breakage at a single site (either *a* or *b*) creates one broken sister chromatid and one nicked sister chromatid, whereas fork breakage at both *a* and *b* creates two broken sister chromatids. (**B)**, Intact *versus* broken sister chromatids were scored and presented as a percentage of the total scored pulse-labeled tracts. Numbers indicate the mean tracts %. Data are from *N*=3 independent experiments.

In contrast to DNA combing, following DNA spreading, ∼90% of labelled fibers contained adjacent red-green tracts, with a much lower percentage of green-only tracts (∼10%) (Fig.4). These data suggest that, as predicted, homologous sister chromatids largely remain adjacent during DNA spreading. Unfortunately, because of the nature of the spreading technique, which results in non-uniform distribution and DNA with frequent crossing, it was not possible to clearly distinguish between red-green-red and red-green tracts.

**Fig. 4.**
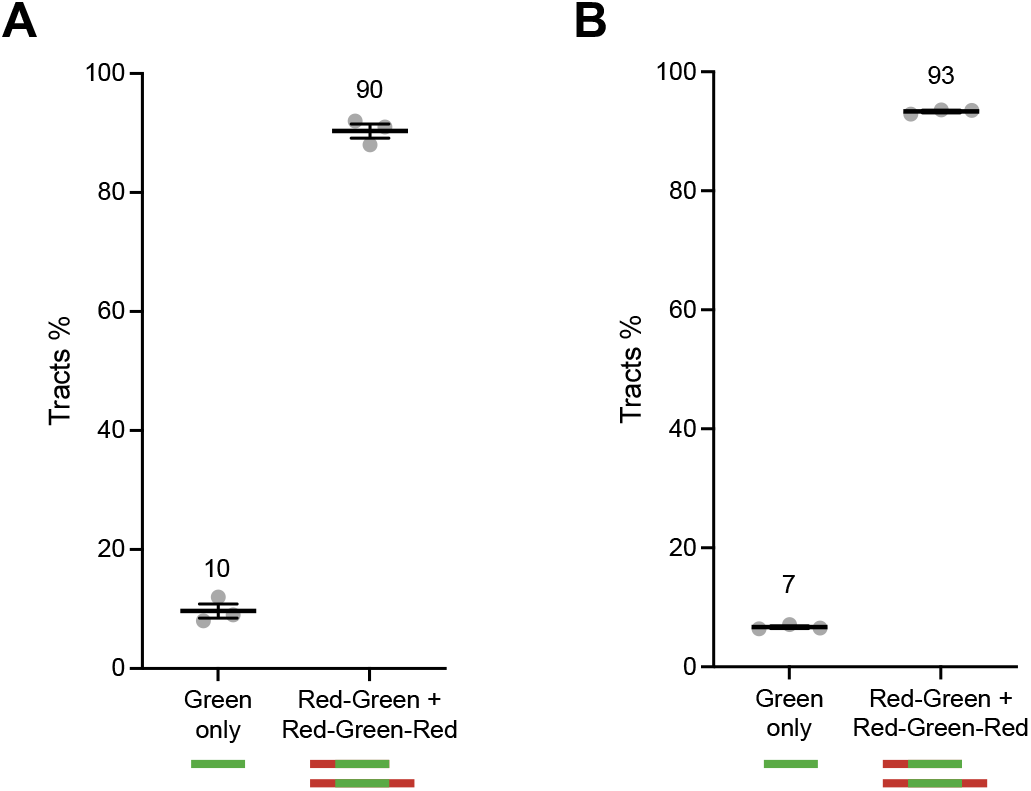
Sister chromatid cohesion is retained during DNA spreading. **(A, B)** DNA spreading assay performed as depicted in Fig. 1A. Green-only *versus* Red/Green tracts were scored and represented as a percentage of total pulse-labelled tracts. Numbers indicate the mean tracts %. **A** and **B** indicate results from the Vindigni and Chaudhuri laboratories, respectively (*N*=3 in **A** and *N*=3 in **B**).

As an additional test of our model, we compared the ability of DNA spreading and DNA combing to detect DNA single-strand gaps (ssDNA gaps). Post-replicative single-strand DNA gaps have emerged as a vulnerability of cancer cells and are detectable by pre-treatment of genomic DNA with the single-strand specific S1 nuclease enzyme prior to DNA spreading or DNA combing (S1 fiber technique) (Quinet et al. 2016; Vaitsiankova et al. 2022). This is because S1 nuclease cleaves the intact DNA strand at ssDNA gaps, resulting in shortening of the DNA replication tract associated with that sister chromatid. Based on our model, S1 treatment should lead to different outcomes in DNA spreading and DNA combing experiments. Specifically, if DNA combing separates sister chromatids as we have proposed, then treatment with S1-nuclease will unveil ssDNA gaps and thus lead to detectable replication tract shortening as described above. In contrast, if DNA spreading does not separate sister chromatids, treatment with S1 nuclease will not lead to detectable shortening of DNA replication tracts because the presence of the gap will be masked by the adjacent sister chromatid, except in the situation of closely apposed ssDNA gaps in both adjacent sister chromatids. To test this idea, we treated cells with an inhibitor of the lagging-strand maturation protein FEN1 nuclease (FEN1i) (Tumey et al. 2005; Exell et al. 2016), prior to measuring the size of DNA replication tracks by DNA spreading or DNA combing. FEN1 nuclease is an enzyme essential for the Okazaki fragment processing (Harrington and Lieber 1994; Goulian et al. 1990), and we have shown previously that FEN1i induces PARP activation and single-strand gaps at unligated Okazaki fragments (Vaitsiankova et al. 2022; Hanzlikova et al. 2018). We labeled RPE-1 cells with CldU for 20 minutes and then with IdU for 60 minutes in presence or absence of FEN1i. Next, we performed the S1 DNA spreading and combing experiment in parallel (Fig 5A). As predicted by our model, DNA spreading was unable to detect S1-dependent shortening in cells treated with FEN1i (Fig 5B). In contrast, we readily detected S1-dependent shortening of DNA replication tracts using DNA combing, confirming that this technique physically separates homologous sister chromatids (Fig 5B).

**Fig. 5.**
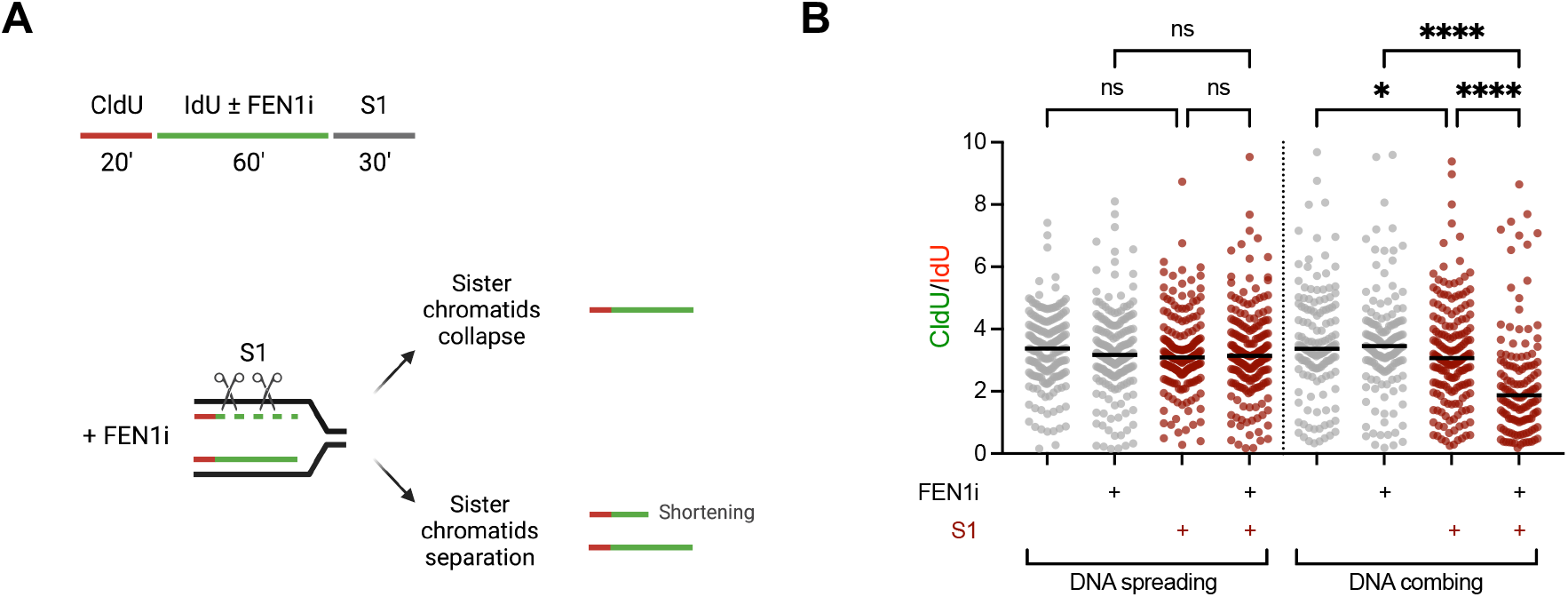
DNA combing, but not DNA spreading, can detect DNA single-strand gaps within individual sister-chromatids. (**A)**, Schematic depicting the use of pulse-labeling and FEN1 inhibitor (FEN1i) to detect single-strand gaps at unligated Okazaki fragments. (**B)**, Dot plots of IdU/CldU ratios in cells treated ± FEN1i and ± S1 nuclease as indicated (*N*=2). Bars represent the median values.

In conclusion, our data demonstrate that the DNA spreading technique does not resolve the two sister chromatids, whereas DNA combing does. Addressing this question is crucial to properly interpret DNA fiber experiments, and to determine whether defects in DNA replication fork progression and/or the maturation of nascent DNA occurs in only the leading or lagging strand, or in both (Tirman et al. 2021; Quinet et al. 2020; Cong et al. 2021; Bai et al. 2020; Mann et al. 2022; Schrempf et al. 2022; Belan et al. 2022; Simoneau, Xiong, and Zou 2021). For example, only the use of DNA combing allows the detection of DNA single-strand gaps and/or DNA double-strand breaks arising in one of two adjacent sister chromatids, resulting for example from unprocessed Okazaki fragments, DNA replication fork bypass of a template strand DNA lesion, or DNA replication fork collision with a DNA single-strand break.

## Materials and Methods

### DNA spreading

The DNA fiber spreading assay was performed in the Vindigni lab, as previously described (Meroni et al. 2022). Exponentially growing RPE-1 (human retinal pigment epithelial) cells were pulse-labeled with CldU (5-Chloro-2’-deoxyuridine, Millipore Sigma) and IdU (5-Iodo-2’-deoxyuridine, Millipore Sigma) as described in the Figure legends. Cells were then washed twice with PBS, collected by trypsinization, resuspended in PBS for a final concentration of 1,500 cells/μL, and spotted onto positively charged glass slides. Cells were mixed with lysis buffer (200 mM Tris-HCl pH 7.5, 50 mM EDTA, 0.5% SDS in water), incubated 5 min at R.T., and slides were tilted at a 20-45° angle to spread the fibers at a constant, low speed. After 10 min air drying, DNA was fixed with a freshly prepared solution of methanol and glacial acetic acid at 3:1 for 5 min. For immuno-staining, DNA was rehydrated in PBS twice for 5 min, then denatured with 2.5 M HCl for 1 h at R.T. Slides were then washed with PBS three times and blocked with 5% BSA at 37°C for 45 min. DNA fibers were immuno-stained with rat anti-BrdU (1/75, Ab6326, Abcam, RRID: AB_305426) and mouse-anti-BrdU (1/20, 347580, BD Biosciences, RRID: AB_400326) for 1.5 h at R.T., washed three times with PBS-0.05%Tween-20 for 5 min, then incubated with anti-rat Alexa Fluor 488 and anti-mouse Alexa Fluor 568 (1/100, A-21470 and A-21124, ThermoFisher Scientific) for 1 h at R.T.

After three washes with PBS-0.05%Tween-20 of 5 min each, slides were mounted with Prolong Gold Antifade Reagent (P36930, ThermoFisher Scientific). Images were acquired with LAS AF software using a Leica DMi8 confocal microscope with 40x/1.15 oil immersion objective. The DNA fiber spreading assay with the ssDNA-specific S1 nuclease was performed as previously described (Meroni et al. 2022). Briefly, after analogs incorporation, cells were permeabilized with CSK100 (100 mM NaCl, 10 mM MOPS pH 7, 3 mM MgCl2, 300 mM sucrose and 0.5% Triton X-100 in water), treated with the S1 nuclease (18001-016, ThermoFisher Scientific) at 20 U/mL in S1 buffer (30 mM sodium acetate pH 4.6, 10 mM zinc acetate, 5% glycerol, 50 mM NaCl in water) for 30 min at 37°C, and collected in PBS-0.1%BSA with cell scraper. Nuclei were then pelleted at ∼4600 x g for 5 min at 4°C, resuspended in PBS, and processed as intact cells in the standard DNA spreading assay.

The DNA spreading assay in the Chaudhuri lab was conducted as follows. RPE-1 (human retinal pigment epithelial) cells were seeded on a 18×24 mm coverslip and the next day they were pulse-labeled with CldU (5-Chloro-2’-deoxyuridine, Millipore Sigma) and IdU (5-Iodo-2’-deoxyuridine, Millipore Sigma) as described in the Figure legends. Cells were then washed twice with PBS and using a combing machine, the cells were lysed and the DNA spread on the coverslip with lysis buffer (200 mM Tris-HCl pH 7.5, 50 mM EDTA, 0.5% SDS in water). After air drying DNA was fixed with a freshly prepared solution of methanol and glacial acetic acid at 3:1 overnight. For immuno-staining, DNA was rehydrated in PBS twice for 5 min, then denatured with 2.5 M HCl for 1 h at R.T. Slides were then washed with PBS three times and blocked with 5% BSA at RT for 45 min. DNA fibers were immuno-stained with rat anti-BrdU (1/100, Ab6326, Abcam, RRID: AB_305426) and mouse-anti-BrdU (1/100, 347580, BD Biosciences, RRID: AB_400326) for 1.5 h at R.T., washed three times with PBS-0.05%Tween-20 for 5 min, then incubated with anti-rat Cy3 (1:250, Jackson Immuno-Reasearch Laboratories, Inc., Cat# 712-166-153) and anti-mouse Alexa Fluor 488 (1:250, Invitrogen, Cat# A11001) for 1 h at R.T. After three washes with PBS-0.05%Tween-20 of 5 min each, slides were mounted with Prolong Gold Antifade Reagent (P36930, ThermoFisher Scientific). Images acquired by Metafer5 at 40x and data analysis was carried out with ImageJ software64. The DNA fiber spreading assay with the ssDNA-specific S1 nuclease was performed as previously described (Meroni et al. 2022).

### DNA combing

In the Vindigni lab, DNA combing was conducted as follows. Exponentially growing RPE-1 cells were pulse-labeled with CldU (5-Chloro-2’-deoxyuridine, Millipore Sigma) and IdU (5-Iodo-2’-deoxyuridine, Millipore Sigma) as described in the text. Cells were then collected by trypsinization and embedded into agarose plugs, and DNA combing was performed according to manufacturer’s protocol (Genomic Vision) with minor modifications, using the Genomic Vision FiberPrep kit and a Genomic Vision combing machine. The DNA was then baked for 2 hours at 60 °C, and stored at -20 °C. For immuno-staining, DNA was denatured with fresh 0.5 M NaOH, 1 M NaCl solution for 8 min at R.T. Coverslips were then washed three times with PBS, dehydrated with 70%, 90%, and 100% ethanol for 2 min each, and blocked with 10% goat serum in PBS-0.1%Tween-20 at R.T. for 1 h. DNA fibers were immuno-stained with rat anti-BrdU (1/75, Ab6326, Abcam, RRID: AB_305426) and mouse-anti-BrdU (1/20, 347580, BD Biosciences, RRID: AB_400326) for 1 h at 37 °C, washed three times with PBS-0.01%Tween-20, then incubated with anti-rat Alexa Fluor 488 and anti-mouse Alexa Fluor 568 (1/100, A-21470 and A-21124, ThermoFisher Scientific) for 45 min at R.T. After three washes with PBS-0.01%Tween-20, coverslips were mounted with Prolong Gold Antifade Reagent (P36930, ThermoFisher Scientific). For total DNA staining, coverslips are incubated with anti-ssDNA (1/100, MAB3034, Millipore) for 1 h at 37 °C, washed, incubated with anti-mouse Alexa Fluor 647 (1/100, A28181, ThermoFisher Scientific) for 45 min at R.T., washed, and then mounted. Images were acquired with LAS AF software using a Leica DMi8 confocal microscope with 40x/1.15 oil immersion objective. The DNA combing assay with the ssDNA-specific S1 nuclease was performed as follows. Immediately before combing, the DNA solution is mixed 1:1 with 2X S1 buffer (60 mM sodium acetate pH 4.6, 20 mM zinc acetate, 10% glycerol, 100 mM NaCl in water) in presence or absence of 40 U/mL of the S1 nuclease (18001-016, ThermoFisher Scientific), and incubated at R.T. for 30 minutes.

DNA combing in the Caldecott lab was conducted essentially as described previously (Vaitsiankova et al. 2022). In brief, exponentially growing RPE-1 cells were pulse-labeled with 1 μM CldU (5-Chloro-2’-deoxyuridine, Merck, C6891) and 250 μM IdU (5-Iodo-2’-deoxyuridine, Merck, I7125) as described in the text. Cells were then collected by trypsinization and resuspended in ice-cold PBS to give a final concentration of 5×10^6^ cells/ml. Of this cell mix, 50 μl was pre-warmed to 50 °C and embedded into an agarose plug (BioRad). The plugs were incubated overnight in 42 °C in proteinase K lysis buffer (2 mg/ml proteinase K, 10 mM Tris-HCl pH 7.5, 100 mM EDTA, 0.5% SDS, 20 mM NaCl).

Next, the DNA plugs were washed two times for 1 h in TE50 buffer (10 mM Tris-HCl pH7.5, 50 mM EDTA, 100 mM NaCl) followed by two times for 1 h in TE buffer (10 mM Tris-HCl pH 7.5, 1 mM EDTA, 100 mM NaCl). The plugs were then melted at 68 °C for 20 min in 1 ml MES (35 mM MES hydrate, 150 mM MES sodium salt, 100 mM NaCl) and cooled to 42 °C for 10 min. The samples were then incubated at 42°C overnight with the addition of 3 μl of β-agarase (NEB). The DNA mix was then gently poured into combing reservoirs containing 1.2 ml MES and the genomic DNA was combed onto salinized coverslips (Genomic Vision) using a combing machine (Genomic Vision) and baked for 2 h at 68 °C. For immuno-staining, DNA was denatured with fresh 0.5 M NaOH 1 M NaCl solution for 8 min at R.T. Coverslips were then washed three times with PBS, dehydrated with 70%, 90%, and 100% ethanol for 1 min each, and blocked with 1% BSA in PBS-0.1%Tween-20 at R.T. for 1 h. DNA fibers were immuno-stained with rat anti-BrdU (1/30, Ab6326, Abcam, RRID: AB_305426) and mouse-anti-BrdU (1/25, 347580, BD Biosciences, RRID: AB_400326) for 1 h at 37 °C, washed three times with PBS-0.01%Tween-20, then incubated with anti-rat Alexa Fluor 568 and anti-mouse Alexa Fluor 488 (1/25, A11077 and A11001, ThermoFisher Scientific) for 45 min at 37 °C. After three washes with PBS-0.01%Tween-20, coverslips were dehydrated in ethanol before mounting on microscope slides with fluoroshield (Merck). the slides were imaged using an Apotome widefield microscope (Zeiss) with ×40 oil objectives. ImageJ software was used to visualize and score the labelled replication tracks.

In the Ray Chaudhuri lab, DNA combing was performed according to manufacturer’s protocol (Genomic Vision) with minor modifications. Exponentially growing RPE-1 cells were pulse-labeled with CldU (5-Chloro-2’-deoxyuridine, Millipore Sigma) and IdU (5-Iodo-2’-deoxyuridine, Millipore Sigma) as described in the Figure legends. Cells were then collected by trypsinization and embedded into agarose plugs. The plugs were incubated in Proteinase K buffer (10 mM Tris pH 7.5, 100 mM EDTA pH 8.0, 1% N-Laurylsarcosine and 1 mg/mL proteinase K) for 48 hours at 37°C. The proteinase K buffer was changed twice during these 48 hours. The plugs were washed several times in TE50 + 100 mM NaCl buffer (10 mM Tris pH 7.5, 50 mM EDTA pH 8.0 and 100 mM NaCl) and stored at 4°C in the dark until DNA extraction. DNA was extracted by washing the plug for one hour with 1x TE pH 7.5 + 500 mM NaCl buffer, after which the plug was washed for 3 times 5 min with MES buffer (50 mM MES hydrate, 50 mM MES sodium salt and 5 M NaCl). The plug in MES buffer was incubated at 68 °C for 30 min after which it was equilibrated at 42 °C for 5 min. 1,2 U β-agarase (NEB, M0392) per mL MES buffer was added to the solution and incubated at 42 °C overnight. The extracted DNA was combed using a combing machine onto silanized coverslips (Genomic Vision), baked for 2 hours at 60 °C, and stored at -20 °C.

For immuno-staining, DNA was denatured with fresh 0.5 M NaOH 1 M NaCl solution for 8 min at R.T. Coverslips were then washed three times with PBS, dehydrated with 70%, 90%, and 100% ethanol for 2 min each, and blocked with 3% BSA in PBS at RT in a humidified chamber for 30 min. DNA fibers were immuno-stained with rat anti-BrdU (1/75, Ab6326, Abcam, RRID: AB_305426) and mouse-anti-BrdU (1/20, 347580, BD Biosciences, RRID: AB_400326) for 45 min at 37 °C in a humidified chamber, washed three times with PBS-0.01%Tween-20, then incubated with anti-rat Cy3 (1:250, Jackson Immuno-Reasearch Laboratories, Inc., Cat# 712-166-153) and anti-mouse Alexa Fluor

488 (1:250, Invitrogen, Cat# A11001) for 45 min at 37 °C in a humidified chamber. After three washes with PBS-0.01%Tween-20, coverslips were mounted with Prolong Gold Antifade Reagent (P36930, ThermoFisher Scientific). For total DNA staining, coverslips are incubated with anti-ssDNA (1/100, MAB3034, Millipore) for 2 h at 37 °C in a humidified chamber, washed, incubated with anti-mouse Alexa Fluor 647 (1/100, A28181, ThermoFisher Scientific) for 45 min at 37 °C in a humidified chamber, washed, and then mounted. Images acquired by Metafer5 at 40x and data analysis was carried out with ImageJ software64. The DNA combing assay with the ssDNA-specific S1 nuclease was performed as follows. Immediately before combing, the DNA solution is mixed 1:1 with 2X S1 buffer (60 mM sodium acetate pH 4.6, 20 mM zinc acetate, 10% glycerol, 100 mM NaCl in water) in presence or absence of 40 U/mL of the S1 nuclease (18001-016, ThermoFisher Scientific), and incubated at R.T. for 30 minutes.

## Acknowledgments

Work in the AV lab was supported by the National Cancer Institute (NCI) grants R01CA237263 and R01CA248526, the U.S. Department of Defense (DOD) Breast Cancer Research Program (BRCP) Expansion Award BC191374, the Alvin J. Siteman Cancer Center Siteman Investment Program (supported by The Foundation for Barnes-Jewish Hospital, Cancer Frontier Fund), and the Barnard Foundation. Work in the Caldecott lab was supported by a Cancer Research UK Programme Grant (C6563/A27322). Work in the Ray Chaudhuri lab was supported by a Dutch Research Council (NWO) VIDI grant (Vidi.193.131), Josephine Nefkens Cancer Program and Ammodo Science Award.

## Author Contributions

KWC/AV/AC conceptualized and developed the project. KWC/SW conceived the idea of the differential labeling scheme (Fig.1, Fig. 2 & Fig. 4), and AM/AV conceived the experiments in Fig. 3 and Fig. 5. AV/KWC/AM wrote the manuscript with help from SW/AC/CF. SW conducted DNA combing in Fig. 2; AM and CF conducted DNA combing and DNA spreading in Fig. 2 & Fig. 3; and AM conducted the DNA breakage and S1 nuclease experiments in Fig. 3 & Fig. 5.

## References

Ashour, M. E., A. K. Byrum, A. Meroni, J. Xia, S. Singh, R. Galletto, S. M. Rosenberg, A. Vindigni, and N. Mosammaparast. 2023. ‘Rapid profiling of DNA replication dynamics using mass spectrometry-based analysis of nascent DNA’, J Cell Biol, 222.

Bai, G., C. Kermi, H. Stoy, C. J. Schiltz, J. Bacal, A. M. Zaino, M. K. Hadden, B. F. Eichman, M. Lopes, and K. A. Cimprich. 2020. ‘HLTF Promotes Fork Reversal, Limiting Replication Stress Resistance and Preventing Multiple Mechanisms of Unrestrained DNA Synthesis’, Mol Cell, 78: 1237–51.e7.

Belan, O., M. Sebald, M. Adamowicz, R. Anand, A. Vancevska, J. Neves, V. Grinkevich, G. Hewitt, S. Segura-Bayona, R. Bellelli, H. M. R. Robinson, G. S. Higgins, G. C. M. Smith, S. C. West, D. S. Rueda, and S. J. Boulton. 2022. ‘POLQ seals postreplicative ssDNA gaps to maintain genome stability in BRCA-deficient cancer cells’, Mol Cell, 82: 4664–80 e9.

Bensimon, A., A. Simon, A. Chiffaudel, V. Croquette, F. Heslot, and D. Bensimon. 1994. ‘Alignment and sensitive detection of DNA by a moving interface’, Science, 265: 2096–8.

Berti, M., D. Cortez, and M. Lopes. 2020. ‘The plasticity of DNA replication forks in response to clinically relevant genotoxic stress’, Nat Rev Mol Cell Biol, 21: 633–51.

Cong, K., M. Peng, A. N. Kousholt, W. T. C. Lee, S. Lee, S. Nayak, J. Krais, P. S. VanderVere-Carozza, K. S. Pawelczak, J. Calvo, N. J. Panzarino, J. J. Turchi, N. Johnson, J. Jonkers, E. Rothenberg, and S. B. Cantor. 2021. ‘Replication gaps are a key determinant of PARP inhibitor synthetic lethality with BRCA deficiency’, Mol Cell, 81: 3227.

Cybulla, E., and A. Vindigni. 2023. ‘Leveraging the replication stress response to optimize cancer therapy’, Nat Rev Cancer, 23: 6–24.

Daigaku, Y., A. A. Davies, and H. D. Ulrich. 2010. ‘Ubiquitin-dependent DNA damage bypass is separable from genome replication’, Nature, 465: 951–5.

Exell, J. C. M. J. Thompson, L. D. Finger, S. J. Shaw, J. Debreczeni, T. A. Ward, C. McWhirter, C. L. Siöberg, D. Martinez Molina, W. M. Abbott, C. D. Jones, J. W. Nissink, S. T. Durant, and J. A. Grasby. 2016. ‘Cellularly active N-hydroxyurea FEN1 inhibitors block substrate entry to the active site’, Nat Chem Biol, 12: 815–21.

Goulian, M., S. H. Richards, C. J. Heard, and B. M. Bigsby. 1990. ‘Discontinuous DNA synthesis by purified mammalian proteins’, J Biol Chem, 265: 18461–71.

Hanzlikova, H., I. Kalasova, A. A. Demin, L. E. Pennicott, Z. Cihlarova, and K. W. Caldecott. 2018. ‘The Importance of Poly(ADP-Ribose) Polymerase as a Sensor of Unligated Okazaki Fragments during DNA Replication’, Mol Cell, 71: 319–31 e3.

Harrington, J. J., and M. R. Lieber. 1994. ‘The characterization of a mammalian DNA structure-specific endonuclease’, EMBO J, 13: 1235–46.

Jackson, D. A., and A. Pombo. 1998. ‘Replicon clusters are stable units of chromosome structure: evidence that nuclear organization contributes to the efficient activation and propagation of S phase in human cells’, J Cell Biol, 140: 1285–95.

Mann, A., M. A. Ramirez-Otero, A. De Antoni, Y. W. Hanthi, V. Sannino, G. Baldi, L. Falbo, A. Schrempf, S. Bernardo, J. Loizou, and V. Costanzo. 2022. ‘POLtheta prevents MRE11-NBS1-CtIP-dependent fork breakage in the absence of BRCA2/RAD51 by filling lagging-strand gaps’, Mol Cell, 82: 4218–31 e8.

Meroni, A., J. Grosser, S. Agashe, N. Ramakrishnan, J. Jackson, P. Verma, L. Baranello, and A. Vindigni. 2022. ‘NEDDylated Cullin 3 mediates the adaptive response to topoisomerase 1 inhibitors’, Sci Adv, 8: eabq0648.

Michalet, X., R. Ekong, F. Fougerousse, S. Rousseaux, C. Schurra, N. Hornigold, M. van Slegtenhorst, J. Wolfe, S. Povey, J. S. Beckmann, and A. Bensimon. 1997. ‘Dynamic molecular combing: stretching the whole human genome for highresolution studies’, Science, 277: 1518–23.

Parra, I., and B. Windle. 1993. ‘High resolution visual mapping of stretched DNA by fluorescent hybridization’, Nat Genet, 5: 17–21.

Quinet, A., D. Carvajal-Maldonado, D. Lemacon, and A. Vindigni. 2017. ‘DNA Fiber Analysis: Mind the Gap!’, Methods Enzymol, 591: 55–82.

Quinet, A., D. J. Martins, A. T. Vessoni, D. Biard, A. Sarasin, A. Stary, and C. F. Menck. 2016. ‘Translesion synthesis mechanisms depend on the nature of DNA damage in UV-irradiated human cells’, Nucleic Acids Res, 44: 5717–31.

Quinet, A., S. Tirman, J. Jackson, S. Svikovic, D. Lemacon, D. Carvajal-Maldonado, D. Gonzalez-Acosta, A. T. Vessoni, E. Cybulla, M. Wood, S. Tavis, L. F. Z. Batista, J. Mendez, J. E. Sale, and A. Vindigni. 2020. ‘PRIMPOL-Mediated Adaptive Response Suppresses Replication Fork Reversal in BRCA-Deficient Cells’, Mol Cell, 77: 461–74 e9.

Schrempf, A., S. Bernardo, E. A. Arasa Verge, M. A. Ramirez Otero, J. Wilson, D. Kirchhofer, G. Timelthaler, A. M. Ambros, A. Kaya, M. Wieder, G. F. Ecker, G. E. Winter, V. Costanzo, and J. I. Loizou. 2022. ‘POLtheta processes ssDNA gaps and promotes replication fork progression in BRCA1-deficient cells’, Cell Rep, 41: 111716.

Simoneau, A., R. Xiong, and L. Zou. 2021. ‘The trans cell cycle effects of PARP inhibitors underlie their selectivity toward BRCA1/2-deficient cells’, Genes Dev, 35: 1271–89.

Techer, H., S. Koundrioukoff, D. Azar, T. Wilhelm, S. Carignon, O. Brison, M. Debatisse, and B. Le Tallec. 2013. ‘Replication dynamics: biases and robustness of DNA fiber analysis’, J Mol Biol, 425: 4845–55.

Tirman, S., A. Quinet, M. Wood, A. Meroni, E. Cybulla, J. Jackson, S. Pegoraro, A. Simoneau, L. Zou, and A. Vindigni. 2021. ‘Temporally distinct post-replicative repair mechanisms fill PRIMPOL-dependent ssDNA gaps in human cells’, Mol Cell, 81: 4026–40 e8.

Tumey, L. Nathan, David Bom, Bayard Huck, Elizabeth Gleason, Jianmin Wang, Daniel Silver, Kurt Brunden, Sherry Boozer, Stephen Rundlett, Bruce Sherf, Steven Murphy, Tom Dent, Christina Leventhal, Andrew Bailey, John Harrington, and Youssef L. Bennani. 2005. ‘The identification and optimization of a N-hydroxy urea series of flap endonuclease 1 inhibitors’, Bioorganic & Medicinal Chemistry Letters, 15: 277–81.

Vaitsiankova, Alina, Kamila Burdova, Margarita Sobol, Amit Gautam, Oldrich Benada, Hana Hanzlikova, and Keith W. Caldecott. 2022. ‘PARP inhibition impedes the maturation of nascent DNA strands during DNA replication’, Nature Structural & Molecular Biology, 29: 329–38.

Yao, N. Y., and M. O’Donnell. 2009. ‘Replisome structure and conformational dynamics underlie fork progression past obstacles’, Curr Opin Cell Biol, 21: 336–43.

